# Cannabidiol effects on cocaine-seeking behaviour and incubation of craving in mice

**DOI:** 10.1101/2020.12.18.423391

**Authors:** Laia Alegre-Zurano, Miguel Á. Luján, Lídia Cantacorps, Ana Martín-Sánchez, Alba García-Baos, Olga Valverde

## Abstract

**Background and Purpose:** To remain abstinent represents one of the major challenges for the treatment of cocaine use disorder. Cocaine seeking elicited by drug-associated cues progressively intensifies during abstinence in a process termed incubation of craving, representing an aggravating factor for relapse. Cannabidiol is a phytocannabinoid that exerts protecting effects upon cocaine-seeking behaviour, although its effects on cocaine-craving incubation have never been elucidated.

**Experimental Approach:** We developed a mouse model of behavioural economic analysis of demand curves and incubation of cue-induced cocaine craving. Changes in the protein expression of AMPAR subunits and ERK_1/2_ phosphorylation were analysed. We also assessed the effects of cannabidiol (20 mg·kg^-1^) administered either during acquisition of cocaine self-administration or abstinence.

**Key Results:** Mice efficiently performed the demand task and incubation of cocaine craving. Besides, changes in GluA1 and GluA2 protein levels were found along the abstinence in prelimbic cortex, ventral striatum and amygdala, as well as a decrease in ERK_1/2_ phosphorylation in ventral striatum. Cannabidiol reduced ongoing cocaine intake when administered during the acquisition phase of the self-administration, but failed to alter the subsequent demand task performance and incubation of cocaine craving. No effects were found when cannabidiol was administered during the abstinence period.

**Conclusion and Implications:** We provide here a novel model of behavioural economic analysis of demand curves and cue-induced incubation of cocaine-seeking behaviour for mice. Moreover, we show that cannabidiol exerts differential effects on the current model depending on the self-administration phase in which it was administered.

**What is already known:**

- Behavioural economics and incubation of cocaine craving are well-stablished paradigms to evaluate cocaine seeking in rats.
- CBD reduces cocaine-seeking and cocaine-taking behaviours.

**What this study adds:**

- A mouse model of behavioural economic analysis of demand curves and incubation of cue-induced cocaine craving.
- CBD reduces cocaine self-administration and has no effect over demand task and cocaine-craving incubation.

**Clinical significance:**

- A new behavioural model for studying cocaine addiction in mice.
- CBD exerts differential effects depending on when it was administered in the addictive process.

**Tables of Links:** 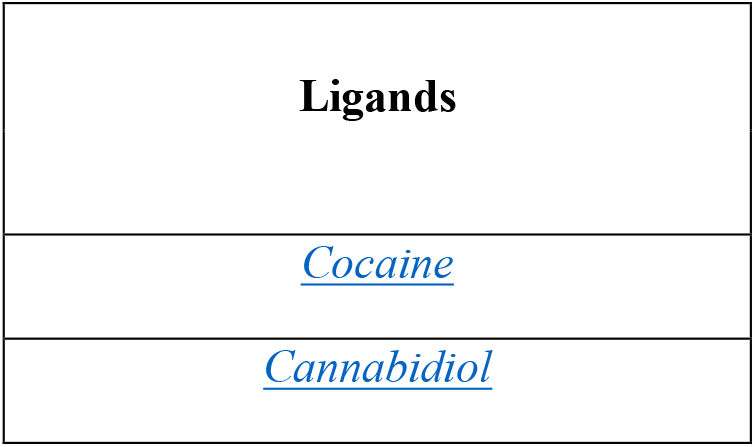

## 1. INTRODUCTION

Cocaine is one of the most used drugs worldwide, showing an increasing trend and doubling the prevalence in Europe compared to the global data (United Nations publication, 2020). However, despite the extended use of cocaine among the population and the problem that it represents for Public Health, the pharmacotherapeutic strategies used to treat cocaine use disorder to date seem to have limited efficacy (Kampman, 2019).

The difficulty to stay abstinent and the risk of relapse represent the major challenges for the treatment of cocaine use disorder. In this sense, cue-induced cocaine craving is one of the main factors contributing to relapse in abstinent addicts. Cocaine-seeking elicited by drug-associated cues progressively intensifies during abstinence in a process termed incubation of craving (Grimm et al., 2001), firstly described in animal models. Clinical studies have already provided evidence about the incubation of craving for cocaine (Parvaz et al., 2016) and other drugs of abuse (Bedi et al., 2011; Wang et al., 2013; Li et al., 2015), although a mechanistic explanation for this phenomenon is yet to be found.

In contrast to the scarce studies in humans, rodent models have been largely used to describe incubation of cocaine craving and to decipher the molecular and cellular mechanisms underlying this process (Lu et al., 2004). Incubation of cocaine craving requires from structural and functional changes in different brain areas, mainly located in the mesocorticolimbic system (Pickens et al., 2011; Wolf, 2016). The accumulation of calcium-permeable α-amino-3-hydroxy-5-methyl-4-isoxazolepropionic acid receptors (CP-AMPARs) lacking GluA2 subunit in the nucleus accumbens (NAc) core is needed for the expression of cocaine-craving incubation after prolonged withdrawal (Conrad et al., 2008). These neuroplastic changes are specially significant since they functionally translate into increases in the number and strength of NAc core neurons responding to cocaine-associated cues (Hollander and Carelli, 2005; Purgianto et al., 2013). Besides, activation of glutamatergic signalling in the medial prefrontal cortex and amygdala are also required for the expression of cocaine-craving incubation. In particular, glutamatergic signalling within the prelimbic cortex (PL) is a key element for the incubation of cocaine-seeking behaviour during late withdrawal (West et al., 2014). More precisely, the glutamatergic PL-NAc core pathway promotes incubation, while the infralimbic cortex (IL)-NAc shell signalling prevents it, based on a process of silent synapses remodelling (Ma et al., 2014). Similarly, distinct nuclei of the amygdala are also differentially implicated in the modulation of drug-seeking behaviour along abstinence. Previous research has demonstrated that both, central nucleus of the amygdala (CeA) and basolateral amygdala (BLA) participate in the first stages of cocaine withdrawal (Lu et al., 2005a). However, their role in prolonged withdrawal is still a matter of debate. Some studies evidence that ERK phosphorylation and glutamatergic activation of CeA, but not BLA, are required for the expression of cocaine-craving incubation during the late withdrawal (Lu et al., 2005b, 2007). Others propose that the glutamatergic BLA-NAc shell pathway is essential for this process (Lee et al., 2013). Furthermore, a recent work identifies the CeA as a common substrate of the incubation of drugs (including cocaine) and natural reinforcers (Roura-Martínez et al., 2019).

Despite the vast research on cocaine-craving incubation in preclinical studies, most of the records to date have been extracted from rat models and only a handful of articles have been published addressing this issue in mouse models (Terrier et al., 2015; Nugent et al., 2017). Besides, long-access self-administration (6h/day) has been the preferred protocol to induce incubation of cocaine craving in both mice and rats (Conrad et al., 2008; Terrier et al., 2015; Calipari et al., 2019).

In the last years, cannabidiol (CBD) has emerged as a new potential treatment for drug abuse (Chye et al., 2019), including cocaine (Calpe-López et al., 2019; Rodrigues et al., 2020). CBD’s multi-target nature (Izzo et al., 2009) seems to contribute to its effectiveness in treating different psychiatric disorders, such as drug addiction. However, it is maybe also the reason why the molecular mechanisms underlying its effects are so complex and, frequently, poorly understood. Although different preclinical studies have evaluated the effects of CBD on acquisition of cocaine self-administration (Mahmud et al., 2016; Luján et al., 2018), extinction (Lujan et al., 2020), reinstatement (Mahmud et al., 2016; Gonzalez-Cuevas et al., 2018; Luján et al., 2018; Lujan et al., 2020), and spontaneous cocaine withdrawal (Gasparyan et al., 2020), its effects on abstinence and cocaine-craving incubation have not yet been confirmed. Of note, CBD or CBD-containing drugs have been clinically tested for the treatment of tobacco (Hindocha et al., 2018) and cannabis (Crippa et al., 2013; Allsop et al., 2014; Trigo et al., 2018) withdrawal, but not for cocaine. Moreover, none of these studies specifically assessed incubation of craving.

In the present study, we implemented a demand task analysed under the behavioural economics paradigm in CD1 male mice exposed to a cocaine-induced self-administration procedure. This strategy allowed us to evaluate relevant parameters related to cocaine-seeking and -taking behaviours. Then, we set up a new procedure for incubation of cue-induced craving in mice during cocaine withdrawal from short-access cocaine self-administration (2h·day^-1^). AMPAR subunit composition and extracellular signal-regulated kinase 1 and 2 (ERK1/2) phosphorylation were determined in PL, ventral striatum (vSTR) and amygdala during the incubation of craving. Finally, we examined the effects of CBD on cocaine-induced acquisition of self-administration demand task and incubation of cocaine craving.

## 2. METHODS

### 2.1. Animals

Male CD1 mice (postnatal day 56) were purchased from Charles River (Barcelona, Spain) and transported to our animal facility (UBIOMEX, PRBB). Animals were maintained in a 12-hour light-dark cycle (lights on 7:30-19:30). Mice were housed at a stable temperature (22 °C ± 2) and humidity (55% ± 10%), with food and water *ad libitum*, and they were allowed to acclimatize to the new environmental conditions for at least five days prior to surgery. Once mice were recovered from surgery, they were moved to the experimental room in an inversed 12-hour light-dark cycle (lights on 19:30-7:30). All animal care and experimental protocols were approved by the UPF/PRBB Animal Ethics Committee (CEEA-PRBB-UPF), in accordance with European Community Council guidelines (2016/63/EU).

### 2.2. Experimental design

#### Experiment 1: Cocaine self-administration, demand task and incubation of craving

Mice were first trained to self-administer cocaine in a fixed ratio (FR) 1 and FR3 schedules of reinforcement, and 24 hours later underwent the demand task. Afterwards, mice started the abstinence period in their home cages and were tested for cue-induced seeking on withdrawal days (WD) 1, 15, 30, and 45 (Figure 1A). Besides, brains were extracted immediately after the cue-induced seeking tests on WD1, WD15 and WD30 for Western blot analysis (Figure 2a).

**Figure 1.**
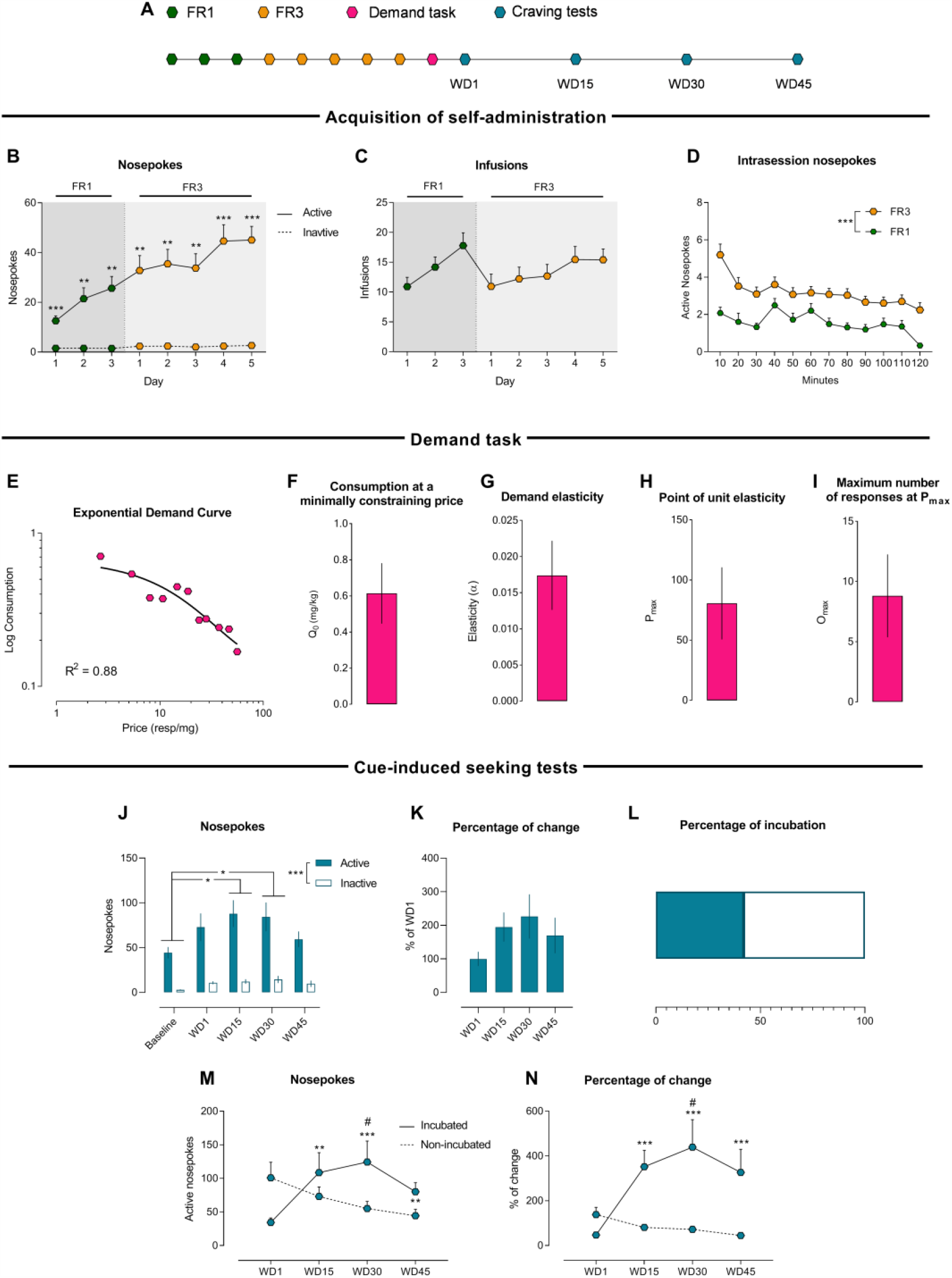
Behavioural economic analysis of demand curve and incubation of cocaine craving paradigms for mice. (A) Schematic timeline of the experiment 1. (B) Active and inactive nosepokes during FR1 and FR3 (n=22) in the self-administration paradigm. Tukey’s test, **p<0.01 and ***p<0.001, active vs inactive nosepokes (C) Infusions during FR1 and FR3. (D) Intrasession active nosepokes during FR1 and FR3. Two-way ANOVA, ***p<0.001, FR1 vs FR3. (E) Exponential demand curve where consumption plotted as a function of price (n=19). (F) Consumption at a minimally constraining price or Q_0_. (G) Demand elasticity or α. (H) Point of unit elasticity or P_max_. (I) Maximum number of responses at P_max_ or O_max_. (J) Active and inactive nosepokes during cue-induced seeking tests (n=19). Two-way ANOVA, ***p<0.001 active vs inactive hole; Tukey’s test, *p<0.01, baseline vs WD15 and WD30. (K) Percentage of active nosepokes change from WD1. (L) Percentage of mice that reached incubation criteria. (M) Active nosepokes of incubated (n=8) and non-incubated (n=11) mice. Sidak’s test, **p<0.01 and ***p<0.001, vs WD1; #p<0.05, vs non-incubated mice. (N) Percentage of active nosepokes change from WD1 of incubated and non-incubated mice. Sidak’s test, ***p<0.001, vs WD1; #p<0.05, vs non-incubated mice.

**Figure 2.**
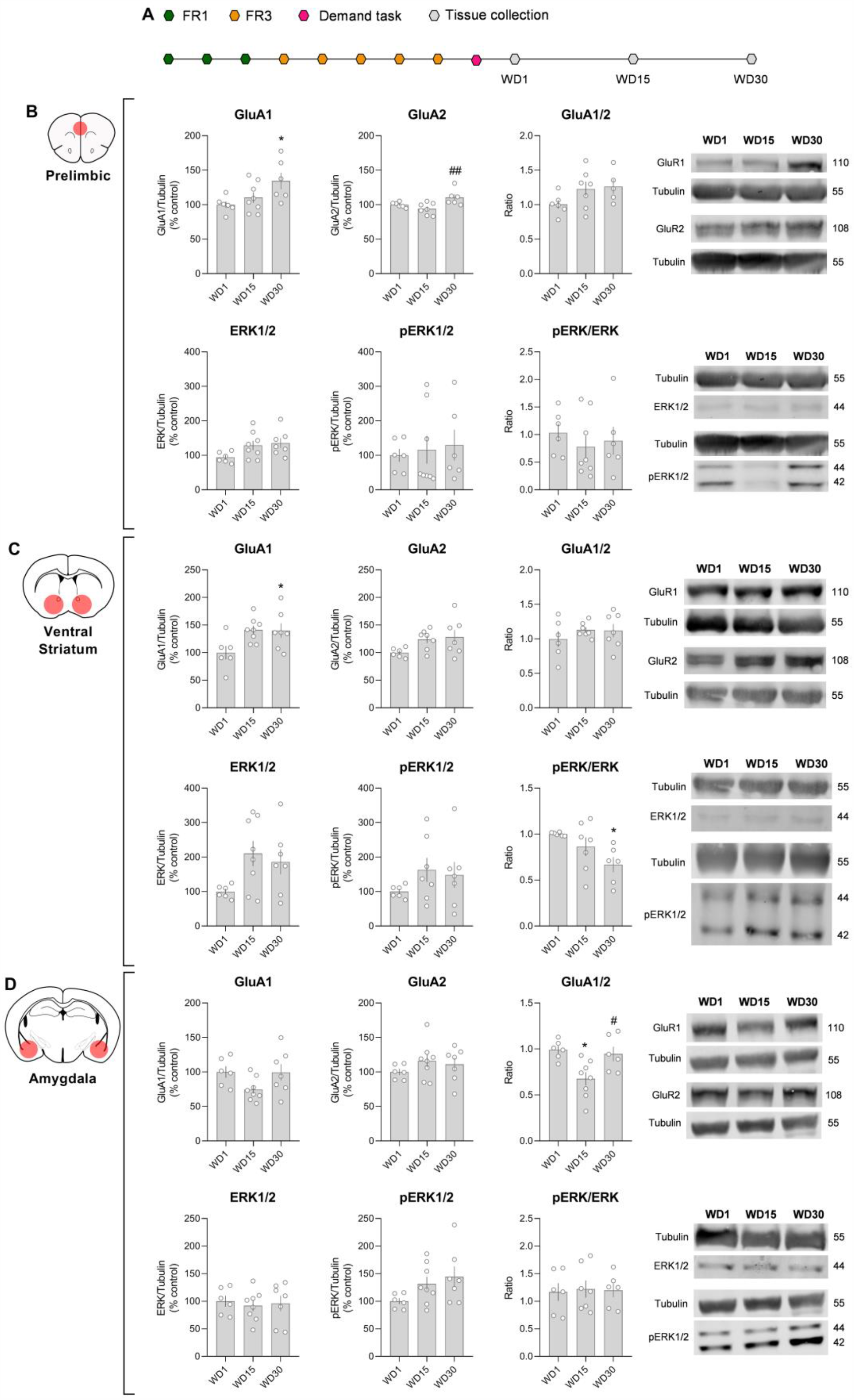
AMPAR subunit composition and ERK phosphorylation changes along abstinence in PL, vSTR and amygdala (n=6-8). (A) Schematic timeline of the experiment 1. GluA1/2 and pERK/ERK changes on WD1, WD15 and WD30 in: (B) PL; Tukey’s test; *p<0.05, vs WD1; ##p<0.01, vs WD15; (C) vSTR; Tukey’s test, *p<0.05, vs WD1; (d) Amygdala; Tukey’s test, *p<0.05, vs WD1; #p<0.05, vs WD15.

#### Experiment 2: Effects of CBD administered during acquisition of cocaine self-administration

Mice followed the same protocol as in Experiment 1, except that vehicle or CBD were administered just before entering the operant conditioning cages during the acquisition of cocaine self-administration and also immediately prior to the demand task. Therefore, as the length of the acquisition phase depends on how quick animals learn the task, the number of days and the number of CBD injections varied between 9 and 16. Mice that succeeded to acquire self-administration behaviour underwent cue-induced seeking tests on WD1 and WD15 (Figure 3a).

**Figure 3.**
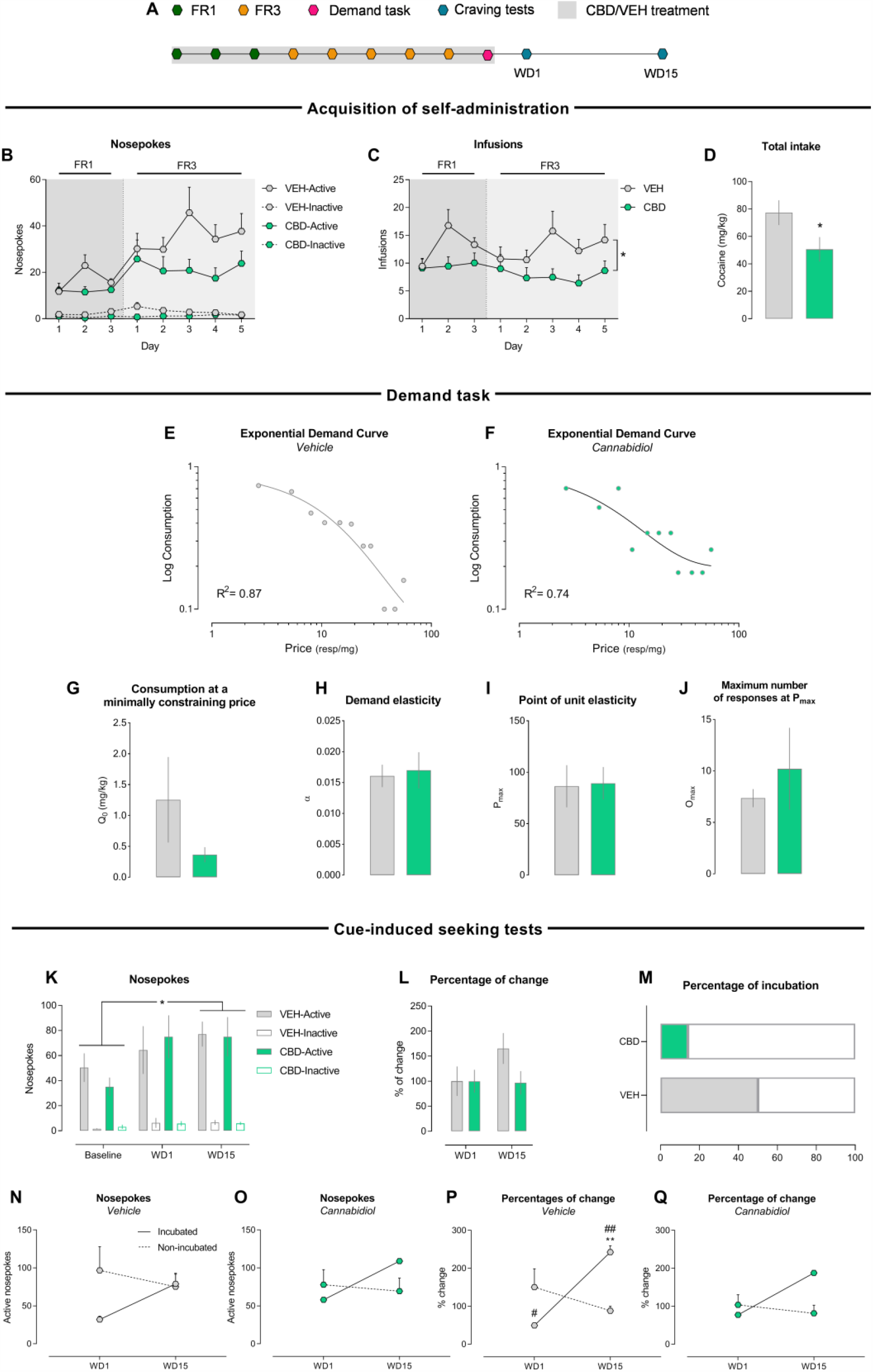
CBD decreases cocaine seeking and taking when administered during acquisition, but such effects are not maintained during abstinence. (A) Schematical timeline of the experiment 2. (B) Active and inactive nosepokes during FR1 and FR3 (Vehicle n=18; CBD n=17). (C) Infusions during FR1 and FR3. Two-way ANOVA, *p<0.05, vehicle- vs CBD-treated group. (D) Total cocaine intake during acquisition of self-administration. Student’s *t* test, *p<0.05. (E) Exponential demand curve where consumption is plotted as a function of price for vehicle- (n=11) and (F) CBD-treated group (n=8). (G) Consumption at a minimally constraining price or Q_0_. (H) Demand elasticity or α. (I) Point of unit elasticity or P_max_. (J) Maximum number of responses at P_max_ or O_max_. (K) Active and inactive nosepokes during cue-induced seeking tests (Vehicle n=8; CBD n=7). Tukey’s test, *p<0.05, baseline vs WD15. (L) Percentage of active nosepokes change from WD1. (M) Percentage of incubation. (N) Active nosepokes for vehicle-treated (incubated mice n=4; non-incubated mice n=4) and (O) CBD-treated (incubated mice n=1; mon-incubated mice n=6) groups. (P) Percentage of change from WD1 for vehicle-treated and (Q) CBD-treated groups. Sidak’s test, **p<0.01, vs WD1; #p<0.05 and ##p<0.01, vs non-incubated mice.

#### Experiment 3: Effects of CBD administered during incubation of cocaine craving

Mice followed the same protocol as in Experiment 1, except that CBD (20 mg·kg^-1^) was administered once a day from WD1 to WD15. Mice underwent cue-induced seeking tests on WD1, WD15 and WD30 (Figure 4a).

**Figure 4.**
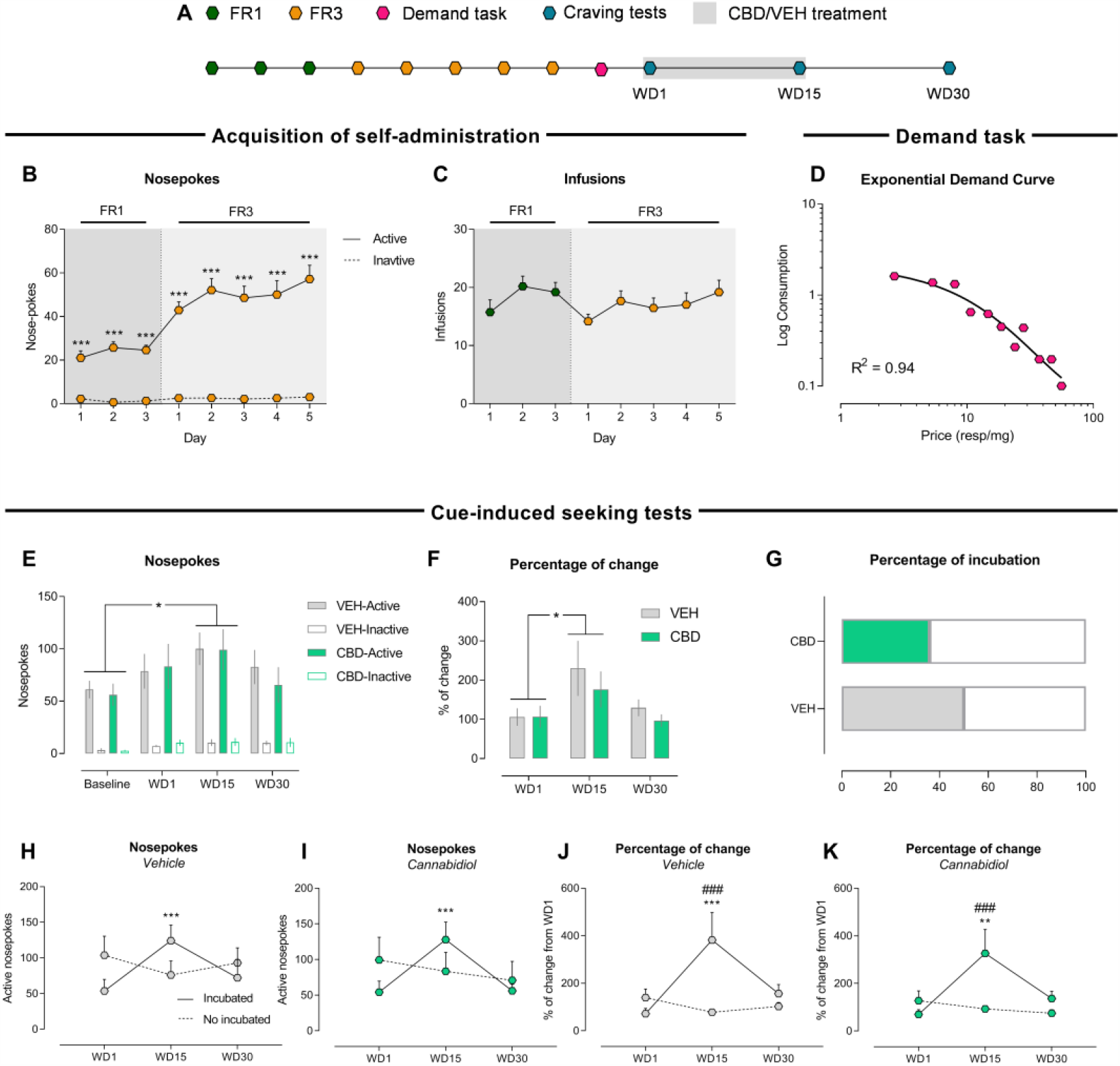
CBD has no effects on cocaine-craving incubation when administered during abstinence. (A) Schematical timeline of the experiment 2. (B) Active and inactive nosepokes during FR1 and FR3 (n=30). Tukey’s test, ***p<0.001 vs inactive hole. (C) Infusions during FR1 and FR3. (D) Exponential demand curve where consumption is plotted as a function of price (n=27). (E) Active and inactive nosepokes during cue-induced seeking tests (Vehicle n=14; CBD n=14). Tukey’s test, *p<0.05, baseline vs WD15. (F) Percentage of active nosepokes change from WD1. Tukey’s test, *p<0.05, WD1 vs WD15. (G) Percentage of mice that reached incubation criteria. (H) Active nosepokes for vehicle-treated (incubated mice n=7; non-incubated mice n=7) and (I) CBD-treated (incubated mice n=5; non-incubated mice n=9) groups. Sidak’s test, ***p<0.001, vs WD1. (J) Percentage of change for vehicle-treated and (K) CBD-treated groups. Sidak’s test, ***p<0.001, vs WD1; Sidak’s test, **p<0.01, vs WD1; ###p<0.001, vs non-incubated mice.

### 2.3. Surgery

The surgical procedure was conducted as previously described (Soria et al., 2005; Lujan et al., 2020) with minor changes. Surgical implantation of the catheter into the jugular vein was performed following anaesthetization with a mixture of ketamine hydrochloride (75 mg·kg^-1^; Imalgène1000, Lyon, France) and medetomidine hydrochloride (1 mg·kg^-1^; Medeson®, Barcelona, Spain). The anaesthetic solution was injected in a volume of 0.15 mL·10 g^-1^ body weight, i.p. Briefly, a 6 cm length of silastic tubing (0.3 mm inner diameter, 0.6 mm outer diameter) (silastic, Dow Corning, Houdeng-Goegnies, Belgium) was fitted to a 22-gauge steel cannula (Semat, Herts, England) that was bent at a right angle and then embedded in a cement disk (Dentalon Plus, Heraeus Kulzer, Germany) with an underlying nylon mesh. The catheter tubing was inserted 1.3 cm into the right jugular vein and anchored with a suture. The remaining tubing ran subcutaneously to the cannula, which exited at the mid-scapular region.

Meloxicam (0.5 mg·kg^-1^ s.c.; Metacam®, Barcelona, Spain), enrofloxacin (7.5 mg·kg^-1^ i.p.; Baytril® 2.5%; Barcelona, Spain), atipamezole hydrochloride (0.5 mg·kg^-1^ i.p.; Revertor®, Barcelona, Spain), and 1 ml glucose 5% solution were injected after the surgery. Home cages were placed on thermal blankets to avoid post-anaesthesia hypothermia. Mice were daily monitored for their weight and treated with meloxicam for 48 h and were allowed to recover for four days before the acquisition phase of the self-administration procedure began.

### 2.4. Acquisition of cocaine self-administration

Cocaine operant self-administration was adapted from Soria et al. (2005) and Luján et al. (2018). The self-administration experiments took place in operant conditioning chambers (Model ENV-307A-CT, Med Associates, Inc. Cibertec. Madrid. Spain) with two nosepoke holes. Animals were connected to a microinfusion pump (PHM-100A, Med-Associates, Georgia, VT, USA). Active and inactive nosepokes were selected randomly. Mice were first trained to self-administer cocaine in a FR1 schedule, where a nosepoke in the active hole resulted in the delivery of a 20 μl cocaine infusion over 2 seconds (0.75 mg·kg^-1^·inf^-1^) and a stimulus light for 4 seconds. Each infusion was followed by a 15-second time-out period. All sessions started with a priming infusion and lasted 2 hours. Mice switched to FR3 schedule for 5 days when the following criteria were met on three consecutive days: a minimum of 5 infusions and 65% of responses received at the active nosepoke. No time-out period was applied in FR3. If animals did not meet the criteria within 10 days under FR1 schedule, they were removed from the study. Mice that failed to increase nosepokes >30% from baseline (three last days of FR1) during 3 out of the 5 days of FR3 were also excluded from the study. No performance-based exclusion criteria were applied in Experiment 2, since the treatments were administered during the acquisition and the fact that some animals failed in acquiring the self-administration behaviour either in FR1 or FR3 was considered an effect of the treatment and, therefore, subject to analysis.

### 2.5. Demand task and behavioural economic analysis

Demand task sessions consisted of a single 2-hour within-session procedure adapted from Bentzley et al. (2013). Acquired mice were placed on the operant cages and received a cocaine priming infusion (0.75 mg·kg^-1^·inf^-1^). The session was divided in 12 10-minute bins, where the requirement for earning an infusion followed an ascending trend on the active nosepoke: 1, 2, 4, 6, 8, 11, 14, 18, 21, 28, 35, 42. Inactive nosepokes had no consequences. The first bin of the session was considered a loading bin where the animal would response in order to achieve the preferred level of intoxication and not in a price-dependent way. Therefore, this first bin was excluded from the analysis. Mice that had no active responses along the session (excluding the loading bin) were excluded from the analysis, since they failed on learning the task.

For the representation and the analysis of Q0 and α, we used a GraphPad template, kindly provided by the Institutes for Behavior Resources (IRB; Baltimore, Maryland, USA). Demand curve was analysed following the exponential model (Hursh and Silberberg, 2008): logQ = logQ_0_ + *k* (*e*^−αQ0C^ − 1). In the exponential demand curve graphs, consumption is plotted as a function of the price (responses·mg^-1^), so when the price increases, the consumption decreases. The value of the constant *k* was automatically calculated for each experiment and set equal for all animals. Q_0_ represents the consumption at a minimally constraining price, meaning the preferred level of consumption with a minimum price. α is a measure of behavioural elasticity, which determines the sensitivity of demand to changes in commodity price (Hursh, 1980). On general terms, a low α value is indicative of little changes in cocaine consumption in response to requirement increases. For the analysis of P_max_ and O_max,_ we used the Excel calculator provided by Kaplan and Reed (2014). P_max_ or point of unit elasticity is considered a metric of motivation, and it refers to the maximal price the animal is willing to pay to maintain the level of consumption. O_max_ is the number of responses that occur at P_max_.

### 2.6. Cue-induced seeking tests

The procedure for cue-induced seeking tests was adapted from Terrier et al. (2015). Mice were placed on the operant cages for 2 hours. An active nosepoke resulted in the presentation of a stimulus light, but no cocaine infusion. Inactive nosepokes had no consequences. An animal was considered to express incubation of craving when the increase in active nosepokes on WD15 was >50% from WD1 and/or the increase on WD15 and 30 was >30% from WD1. On Experiment 2, since cue-induced seeking tests were only undertaken on WD1 and 15, an animal was considered to express incubation of craving when the increase of active nosepokes on WD15 was >50% from WD1.

### 2.7. Western blot

Animals were euthanized by cervical dislocation and PL, vSTR and amygdala were dissected and stored at -80°C. The tissue was homogenized in lysis buffer [0.15M NaCl, 1% TX-100, 10% glycerol, 1mM EDTA, 50mM TRIS pH=7.4 and a phosphatase and protease inhibitor cocktail (Roche, Basel, Switzerland)]. Equal amounts of protein (16 μg) for each sample were mixed with loading buffer (153 mM TRIS pH = 6.8, 7.5% SDS, 40% glycerol, 5 mM EDTA, 12.5% 2-β-mercaptoethanol, and 0.025% bromophenol blue), loaded onto 10% polyacrylamide gels, and then transferred to PVDF sheets (Immobilion-P, MERCK, Burlington, USA). Membranes were blocked for 30 minutes with 5% bovine serum albumin at room temperature and then immunoblotted overnight at 4 °C using primary antibodies. On the next day, membranes were incubated for 1 h with their respective secondary fluorescent antibodies. Protein expression was quantified using an Amersham™ Typhoon™ scanner and quantified using Image Studio Lite software v5.2 (LICOR, USA). Protein expression signals were normalized to the detection of housekeeping control protein in the same samples and expressed in terms of fold-change with respect to WD1 values.

### 2.8. Data and statistical analysis

The data and statistical analysis comply with the recommendations of the British Journal of Pharmacology on experimental design and analysis in pharmacology (Curtis et al., 2018). Except for demand curves, data are presented as mean ± SEM. For statistical analysis we used GraphPad Prism 8.0. software. When one, two or three factors were analysed, we used one, two or three-way ANOVA, respectively. When an experimental condition followed a within-subject design, repeated measures ANOVA was used. The factors used in this study were *schedule* (FR1/FR3), *time* (days or minutes), *hole* (active/inactive), *treatment* (CBD/vehicle) and *incubation* (incubated/non-incubated). The statistical threshold for significance was set at p<0.05. When F achieved significance and there was no significant variance in homogeneity, Tukey’s or Sidak’s *post hoc* (when most appropriate) tests were run. We analysed the results of single factor with two levels, parametric measures with unpaired Student’s *t* tests. Percentage of incubation was analysed using Fisher’s exact test.

### 2.9. Materials

Cocaine HCl (0.75 mg·kg^-1^·inf^-1^) was purchased in Alcaliber S.A. (Madrid, Spain), and was dissolved in 0.9% NaCl. CBD (20 mg·kg^-1^ i.p.) was provided by courtesy of Phytoplant Research S.L., (Córdoba, Spain) and was dissolved in ethanol, cremophor E.L. and 0.9% NaCl (0.5:1:18.5).

For Western blot, the primary antibodies used were anti-ERK_1/2_ (1:1000; Abcam; #ab54230; RRID:AB_2139967), anti-pERK_1/2_ (1:2500; Abcam; #ab50011; RRID:AB_1603684), anti-GluA1 (1:1000; Abcam; ABN241; RRID:AB_2721164), anti-GluA2 (1:1000; Abcam; AB1768-I; RRID:AB_2313802) and anti-beta III Tubulin (1:2500; Abcam; ab18207; RRID:AB_444319). The fluorescent secondary antibodies used were anti-mouse (1:2500; Abcam; #ab216772; RRID:AB_2857338) and anti-rabbit (1:2500; Rockland; 611-144-002-0.5; RRID:AB_11182462).

## RESULTS

### Experiment 1: Mice efficiently perform the demand task and incubation of cocaine craving paradigms

In order to validate the behavioural paradigm under our experimental conditions, a naive group of mice acquired cocaine self-administration, and then underwent demand task and cue-induced seeking tests, as shown on Figure 1A. Regarding the acquisition phase, the factors *time, hole* and their interaction reached significance indicating how mice learned to discriminate between active and inactive hole along the days, showing even a higher number of active responses when they were moved to FR3 (Figure 1B) in order to maintain the number of infusions (Figure 1C). Likewise, the intrasession data showed a significant effect for *time* and *schedule*, but not an interaction, indicating that mice on FR3 performed more active responses than on FR1 throughout the session (Figure 1D).

On demand task, three animals were excluded from the analysis, since they failed to achieve any infusion. The exponential model showed a R^2^ = 0.88 (Figure 1E), suggesting a good fitting of the analysed data. The consumption at a minimally constraining price (Q_0_) was 0.61 mg·kg^-1^ (Figure 1F), demand elasticity (α) was 0.017 (Figure 1F), animals displayed a maximal price (P_max_) to maintain the consumption at 80.65 responses·mg^-1^ (Figure 1H), and the number of responses at that price (O_max_) was 8.81 responses (Figure 1I).

Regarding the cue-induced seeking behaviour, the factors *day* and *hole*, but not their interaction, reached significance, indicating both, discrimination between active and inactive holes and differential responding along withdrawal days (Figure 1J). More precisely, Tukey’s *post hoc* analysis showed that on WD15 and WD30 mice displayed more nosepokes than on baseline. When data were shown as the percentage of active nosepokes change from WD1, no effect was found (Figure 1K).

Finally, mice were divided into two groups depending on whether they exhibited or not craving incubation, according to their performance on the cue-induced seeking tests. When expressing the data as active nosepokes (Figure 1M), no effect was found for *incubation* or *day*, but it was for their interaction. In particular, Sidak’s *post hoc* analysis revealed that incubated mice had more active responses on WD15 and WD30 compared to WD1, and the group that did not express incubation had fewer active responses on WD45 compared to WD1. Besides, incubated mice significantly had more active responses than non-incubated mice on WD30. However, when expressing data as percentage of change from WD1 (Figure 1N), the factors *incubation, day* and their interaction became significant. Sidak’s *post hoc* analysis showed that the group showing incubation increased the percentage of responding on WD15, WD30 and WD45 compared to WD1, while the group that did not incubate showed no differences along days. Also, the percentage of change on WD30 was significantly higher on incubated mice compared to those that did not incubate.

### Experiment 1: GluA1/GluA2 and pERK/ERK proteins levels are modulated during cocaine-craving incubation

To analyse the protein levels of GluA1, GluA2, ERK_1/2_ and pERK_1/2_ during abstinence, mice were euthanised on WD1, WD15 and WD30 immadiately after the cue-induced seeking tests (Figure 2A). Regarding the results obtained in PL (Figure 2B), GluA1 protein expression was increased in WD30 compared to WD1, while GluA2 was increased on WD30 compared to WD15. However, no significant changes on the ratio GluA1/2 were observed. By contrast, both ERK_1/2_ and pERK_1/2_ remained unaltered during the abstinence.

On vSTR (Figure 1C), GluA1 protein levels were increased on WD30 compared to WD1, while no changes in GluA2 or GluA1/2 were observed. Besides, we also found a decrease in the ratio pERK_1/2_ /ERK_1/2_ on WD30 compared to WD1.

Finally, the results obtained in amygdala (Figure 2D) showed a decreased in GluA1/2 ratio on WD15 compared to WD1 and WD30, but no changes in ERK_1/2_ phosphorylation.

### Experiment 2: CBD decreases cocaine seeking and taking when administered during acquisition, but such effects are not maintained during abstinence

CBD treatment was administered just before the acquisition and demand task sessions (Figure 3A). The nosepokes curve shows significant effects for *time, hole* and *treatment*, as well as the interaction between *time* and *hole* (Figure 3B). Regarding infusions, CBD-treated mice significantly self-administered fewer infusions along the acquisition (Figure 3C), as it is also reflected in the total intake (Figure 3D).

For the demand task, mice that failed to meet the acquisition criteria on FR1 and FR3 were excluded from the analyses as they could not meet the requirements of the task. Besides, 3 and 4 mice in the vehicle- and CBD-treated groups, respectively, were also excluded from the analysis as they failed to achieve any infusion. The behavioural economic analysis of the exponential demand curves for vehicle- (Figure 3E) and CBD-treated groups (Figure 3F) did not show significant differences among the different parameters (Figure 3G-J).

In this experiment, we also analysed the effects of CBD treatment administered during the acquisition on the posterior incubation of cocaine craving on WD1 and WD15. The performance during the cue-induced seeking tests revealed a significant effect of *time* and *hole*, but not *treatment* or any interaction, indicating that animals incubated their response regardless of the treatment (Figure 3K). Specifically, we found an increase in the responding on WD15 compared to the baseline. When expressing active nosepokes as percentage of change from WD1, no significant effects were found (Figure 3L). Fisher’s test for percentage of incubation showed no significant differences (Figure 3M).

Finally, vehicle- and CBD-treated mice were divided into incubated or non-incubated groups depending on whether active nosepokes on WD15 showed an increase of >50% from WD1. For the vehicle-treated group, the two-way ANOVA for active nosepokes revealed a significant interaction between *time* and *incubation*, but Sidak’s *post hoc* showed no significant effects (Figure 3N). However, when expressed as percentage of change, incubated mice showed a significant increase in the percentage of change on WD15 while non-incubated group did not (Figure 3P). Besides, Sidak’s *post hoc* analysis also showed that both groups of mice were significantly different on both days. For the CBD-treated group, no significant effects were found when data was expressed as active nosepokes (Figure 3O) or when expressed as percentage of change (Figure 3Q), due to the fact that only one mouse met the criteria for incubation.

### Experiment 3: CBD has no effects on cocaine-craving incubation when administered during abstinence

After mice underwent acquisition for self-administration (Figure 4B, C) and demand task (Figure 4D), they were treated with CBD or vehicle for 15 days and tested for cue-induced seeking on WD1, WD15 and WD30 (Figure 4A). Three-way ANOVA for nosepokes during the cue-induced seeking behaviours revealed a significant effect for *time* and *hole*, but not for *treatment* or any interaction (Figure 4E). More precisely, Tukey’s *post hoc* showed that mice conducted more responses on WD15 compared to the baseline. When data was expressed as percentage of change from WD1, *time*, but not *treatment* nor their interaction, revealed a significant effect, being the percentage of change during WD15 increased compared to WD1 (Figure 4F). Fisher’s test for percentage of incubation showed no significant differences (Figure 4G).

Mice were then divided into two groups depending on their craving-incubation exhibition. For the vehicle-treated group, the two-way ANOVA revealed a significant interaction of *time* and *incubation* for both active nosepokes (Figure 4H) and percentage of change (Figure 4J). For active nosepokes, Sidak’s *post hoc* analysis showed that incubated mice conducted more active responses on WD15 compared to WD1. When expressed as percentage of change, we observed that mice that showed incubation also conducted more active responses on WD15 compared to WD1 and compared to the mice that did not show incubation. For the CBD-treated group, two-way ANOVA for active nosepokes revealed a significant effect for time and the interaction of *time* and *incubation* (Figure 4I). Sidak’s *post hoc* analysis showed that incubated mice performed more active nosepokes on WD15 compared to WD1. When data were expressed as percentage of change from WD1, two-way ANOVA showed a significant effect for *time, incubation* and the interaction between factors (Figure 4K). The *post hoc* analysis revealed that incubated mice increased the percentage of change on WD15 compared to WD1 and to mice that did not show incubation.

## DISCUSION

Incubation of cocaine craving is an important process contributing to the high risk of relapse in both humans and rodents. Mainly, rat models of cue-induced cocaine craving have been used to study the neurobiological components underlying such process. In the present study, we developed a mouse model for behavioural economic analysis of demand curves and incubation of cocaine craving in mice. The molecular analysis indicated changes in GluA1 and GluA2 protein levels along abstinence in PL, vSTR and amygdala, as well as a decrease in ERK_1/2_ phosphorylation in vSTR. Besides, we observed a differential effect of CBD depending on the treatment regime. Thus, CBD was able to reduce cocaine seeking in the self-administration paradigm when administered during the acquisition, but failed to alter the subsequent incubation of cocaine craving. Similarly, CBD administered during the cocaine abstinence period did not exert any effect upon incubation.

Behavioural economic analysis of demand curves has been lately applied to rat models of drug consumption (Bentzley et al., 2014; Calipari et al., 2018). However, this analysis has never been employed in mice despite their extended use in the field of addiction research. Based on previous research (Bentzley et al., 2013), we used a single within-session procedure with ascending unit prices. The increase of the prices in every bin was adapted to make it more suitable for mouse demands, that is, lower increase in each bin compared to rat models. In Experiments 1 and 3, the exponential demand curves had a reasonable fit to the mouse performance on the task (r^2^ 0.88 and 0.94, respectively) and most of the animals were able to complete it consuming at any bin. For Experiment 2, the fit of the model was not as good as expected. This may be due to different reasons. First, since mice were divided into two different groups (vehicle- and CBD-treated mice), the sample size was smaller compared to the previous experiment, complicating the extraction of an accurate curve. Second, this kind of task requires from animals considerably engaged with drug-seeking behaviour. Since administration of CBD during acquisition diminished this behaviour, CBD-treated mice consumed less during the demand task. Thus, the exponential model is provided with more limited information, resulting in a poorer fitting of the data. Keeping these limitations in mind, behavioural analysis of demand curves is a highly valuable tool in addiction research as it offers a wide range of measures, such as consummatory behaviour, motivation and behavioural elasticity (Hursh and Silberberg, 2008; Bentzley et al., 2013), which is now prepared to be applied in mouse models of cocaine self-administration.

We also presented a mouse model for incubation of cocaine craving after short-access cocaine self-administration. To our knowledge, only two studies have previously utilized mice as experimental subjects (Terrier et al., 2015; Nugent et al., 2017). Our findings indicate that WD15 represented the peak of incubation, followed by WD30, whereas on WD45 responding levels were already lower. Similarly, Nugent et al. (2017) found the maximal responding on WD28 also after a short-access cocaine self-administration. Although, in this case, the cocaine dose was lower (0.5 mg·kg^-1^·inf^-1^) than in our study (0.75 mg·kg^-1^·inf^-1^). Furthermore, Terrier et al. (2015) found no craving incubation when applying a short-access self-administration with the same dose used in our study (0.75 mg·kg^-1^·inf^-1^). Nevertheless, since we found the maximal response on WD15 and they only tested cocaine craving on WD1 and WD30, it is possible that the incubation did in fact occur, but they did not observe it.

Regarding the molecular results, we found an increase in GluA1 subunit on WD30 within the vSTR, although the GluA1/2 ratio remained unaltered. Since NAc core and shell have opposite roles on either promoting or inhibiting incubation of cocaine craving, respectively, and we analysed the whole vSTR, it is possible that discrete changes in any of these areas were mutually concealed. Otherwise, larger expositions to the drug (i.e., long access) would possibly cause more evident changes in the glutamatergic system (Purgianto et al., 2013). Surprisingly, Western blot analysis revealed a decrease in ERK_1/2_ phosphorylation on WD30 coinciding with the GluA1 subunit increase. Since ERK_1/2_ phosphorylation is related to neural activity and plasticity, these results could be somehow conflicting. Nevertheless, a study found that ERK_1_ knockout mice presented an enhanced synaptic plasticity in the STR (Mazzucchelli et al., 2002). In the same line, Sun et al. (2015) reported a decrease of pERK_1/2_ in the NAc associated with the incubation of heroin seeking. Otherwise, our results show an increase of both GluA1 and GluA2 protein levels on WD30 in PL, but no differences on the ratio. In accordance, Ma et al. (2014) demonstrated that PL neurons projecting to NAc mature during withdrawal by recruiting non-CP-AMPARs, which could explain the lack of increase in GluA1/2 ratio. Alternatively, albeit both GluA1 and GluA2 subunits are increased on WD30, they could differ in the number of subunits that are either internalised or expressed on the cell surface (Conrad et al., 2008). Besides, we found no changes on ERK_1/2_ phosphorylation along abstinence. Accordingly, Whitfield et al. (2011) also reported no differences in ERK_1/2_ phosphorylation on WD7 in the dorsomedial PFC in rats. Finally, we found a decrease of GluA1/2 ratio on WD15 compared to WD1 and WD30 in the amygdala, although we cannot distinguish whether this finding is due to changes in the CeA or in the BLA, which could imply different functional changes (Wolf, 2016). Furthermore, despite the fact that ERK activation in CeA has been shown to be critical for the incubation of cocaine craving (Lu et al., 2005b), we found no differences throughout abstinence.

Previous work in our laboratory has already provided evidence supporting the beneficial role of CBD in the treatment of cocaine-seeking behaviour. In particular, Luján et al. (2018) demonstrated that CBD (20 mg·kg^-1^), when administered during the acquisition phase (FR1) of cocaine self-administration, is able to reduce cocaine-seeking and -taking behaviours, an effect that is dependent on adult hippocampal neurogenesis (Luján et al., 2019). Similarly, we also found a decrease in cocaine self-administration when CBD was concomitantly administered during the acquisition process. Indeed, such effect was also maintained when the requirements for each infusion became more exigent (FR3). However, CBD-induced decrease in cocaine-seeking behaviour faded during cue-induced cocaine-seeking tests, and both vehicle- and CBD-treated mice incubated their responsiveness over the withdrawal period.

When CBD was administered in the abstinence period, no effect was found during acute (WD1) or prolonged (WD15) treatment, or later (WD30). In a previous study (Gonzalez-Cuevas et al., 2018), CBD was able to reduce drug-seeking behaviour induced by stress or context exposure, although no analysis for incubation of cocaine craving was performed. Moreover, various methodological differences might hinder the comparisons, including the experimental designs, animal species (rat vs mouse), doses of cocaine (0.25 vs 0.75 mg·kg^-1^·inf^-1^) and CBD (15 vs 20 mg·kg^-1^) or routes of administration (transdermal vs intraperitoneal). A more recent study by Gasparyan et al. (2020) demonstrated that CBD normalized motor and somatic disturbances in a spontaneous cocaine withdrawal model in CD1 mice. In this case, CBD would play a beneficial role in the management of physical symptoms associated with cocaine withdrawal, an approach which is substantially different from ours.

In the present study, we applied a criterion for determining whether mice had incubated their seeking behaviour due to the exposition to cocaine-associated cues. Therefore, we can observe considerably different curves for mice that express cocaine-craving incubation and mice that do not. This observation is helpful at the time of validating the following findings: (I) the existence of two different populations of mice regarding their expression of craving incubation, and (II) the lack of CBD effectivity on the incubation of cue-induced cocaine-seeking behaviours, since the curves of both vehicle- and CBD-treated groups resemble the same behaviour.

Altogether, we suggest that CBD only exerts its protecting effects on cocaine-seeking behaviour while learning processes are taking place. Thus, CBD was able to reduce drug-seeking behaviour, but could not influence the formation of long-lasting memories associated to the operant task. Hence, once the treatment ceased, the effects of CBD faded and CBD-treated animals were able to incubate cue-induced seeking behaviour as much as control animals. On the same note, CBD administered during the withdrawal period (once the acquisition of self-administration had finished) was not able to revert the seeking behaviour that had already been established in these animals. Therefore, according to this hypothesis, CBD’s lack of effects could be due to the fact that it was not being concomitantly administered when the operant learning was taking place.

In conclusion, we provide a novel model of behavioural economic analysis of demand curves and cue-induced incubation of cocaine-seeking behaviour in mice. Furthermore, we also provide evidence supporting that CBD is able to reduce cocaine consumption when administered during the acquisition of self-administration but it shows no effects over incubation of cocaine craving, suggesting that CBD is only effective when administered while learning of seeking and taking behaviours are taking place.

## ABBREVIATIONS

BLA: Basolateral amygdala
CBD: Cannabidiol
CeA: Central nucleus of the amygdala
CP-AMPAR: Calcium-permeable α-amino-3-hydroxy-5-methyl-4-isoxazolepropionic acid receptor
ERK_1/2_: Extracellular signal-regulated kinase 1 and 2
R: Fixed ratio
IL: Infralimbic cortex
NAc: Nucleus accumbens
pERK_1/2_: Phosphorylated extracellular signal-regulated kinase 1 and 2 phosphorylation
PL: Prelimbic cortex
vSTR: Ventral striatum
WD: Withdrawal day

## ACKNOWLEDGEMENTS

This study was supported by the Ministerio de Economia y Competitividad (grant number PID2019-104077RB-100), Ministerio de Sanidad (Retic-ISCIII, RD16/017/010 and Plan Nacional sobre Drogas 2018/007). L.A.Z received a FPI grant (BES-2017-080066) from Ministerio de Economia y Competitividad. A.G.B received a FI-AGAUR grant from the Generalitat de Catalunya (2019FI_B0081). The Department of Experimental and Health Sciences (UPF) is a “Unidad de Excelencia María de Maeztu” funded by the AEI (CEX2018-000792-M).

## AUTHOR CONTRIBUTIONS

L.A.Z. and O.V. were responsible for the study concept and design. L.A.Z., L.C, A.M.S. and A.G.B. carried out the experimental studies. L.A.Z. and M.A.L. programmed the demand task code and analysis. L.A.Z and O.V. drafted the manuscript. All authors critically reviewed the content and approved the final version for publication.

## CONFLICT OF INTEREST

The authors declare no conflicts of interest

## DECLARATION OF TRANSPARENCY AND SCIENTIFIC RIGOUR

This Declaration acknowledges that this paper adheres to the principles for transparent reporting and scientific rigour of preclinical research as stated in the BJP guidelines for Design & Analysis, Immunoblotting and Immunochemistry, and Animal Experimentation, and as recommended by funding agencies, publishers, and other organizations engaged with supporting research.

## Notes

### Competing Interest Statement

The authors have declared no competing interest.

## REFERENCES

Allsop, D.J., Copeland, J., Lintzeris, N., Dunlop, A.J., Montebello, M., Sadler, C., et al. (2014). Nabiximols as an Agonist Replacement Therapy During Cannabis Withdrawal. JAMA Psychiatry 71: 281.

Bedi, G., Preston, K.L., Epstein, D.H., Heishman, S.J., Marrone, G.F., Shaham, Y., et al. (2011). Incubation of cue-induced cigarette craving during abstinence in human smokers. Biol. Psychiatry 69: 708–711.

Bentzley, B.S., Fender, K.M., and Aston-Jones, G. (2013). The behavioral economics of drug self-administration: a review and new analytical approach for within-session procedures. Psychopharmacology (Berl). 226: 113–25.

Bentzley, B.S., Jhou, T.C., and Aston-Jones, G. (2014). Economic demand predicts addiction-like behavior and therapeutic efficacy of oxytocin in the rat. Proc. Natl. Acad. Sci. U. S. A. 111: 11822–7.

Calipari, E.S., Godino, A., Peck, E.G., Salery, M., Mervosh, N.L., Landry, J.A., et al. (2018). Granulocyte-colony stimulating factor controls neural and behavioral plasticity in response to cocaine. Nat. Commun. 9: 9.

Calipari, E.S., Godino, A., Salery, M., Damez-Werno, D.M., Cahill, M.E., Werner, C.T., et al. (2019). Synaptic Microtubule-Associated Protein EB3 and SRC Phosphorylation Mediate Structural and Behavioral Adaptations During Withdrawal From Cocaine Self-Administration. J. Neurosci. 39: 5634–5646.

Calpe-López, C., Pilar García-Pardo, M., and Aguilar, M.A. (2019). Cannabidiol treatment might promote resilience to cocaine and methamphetamine use disorders: A review of possible mechanisms. Molecules 24:.

Chye, Y., Christensen, E., Solowij, N., and Yücel, M. (2019). The Endocannabinoid System and Cannabidiol’s Promise for the Treatment of Substance Use Disorder. Front. Psychiatry 10: 63.

Conrad, K.L., Tseng, K.Y., Uejima, J.L., Reimers, J.M., Heng, L.-J., Shaham, Y., et al. (2008). Formation of accumbens GluR2-lacking AMPA receptors mediates incubation of cocaine craving. Nature 454: 118–121.

Crippa, J.A.S., Hallak, J.E.C., Machado-de-Sousa, J.P., Queiroz, R.H.C., Bergamaschi, M., Chagas, M.H.N., et al. (2013). Cannabidiol for the treatment of cannabis withdrawal syndrome: a case report. J. Clin. Pharm. Ther. 38: 162–4.

Curtis, M.J., Alexander, S., Cirino, G., Docherty, J.R., George, C.H., Giembycz, M.A., et al. (2018). Experimental design and analysis and their reporting II: updated and simplified guidance for authors and peer reviewers. Br. J. Pharmacol. 175: 987–993.

Gasparyan, A., Navarrete, F., Rodríguez-Arias, M., Miñarro, J., and Manzanares, J. (2020). Cannabidiol Modulates Behavioural and Gene Expression Alterations Induced by Spontaneous Cocaine Withdrawal. Neurotherapeutics 1–9.

Gonzalez-Cuevas, G., Martin-Fardon, R., Kerr, T.M., Stouffer, D.G., Parsons, L.H., Hammell, D.C., et al. (2018). Unique treatment potential of cannabidiol for the prevention of relapse to drug use: preclinical proof of principle. Neuropsychopharmacology 43: 2036–2045.

Grimm, J.W., Hope, B.T., Wise, R.A., and Shaham, Y. (2001). Neuroadaptation. Incubation of cocaine craving after withdrawal. Nature 412: 141–2.

Hindocha, C., Freeman, T.P., Grabski, M., Stroud, J.B., Crudgington, H., Davies, A.C., et al. (2018). Cannabidiol reverses attentional bias to cigarette cues in a human experimental model of tobacco withdrawal. 113: 1696–1705.

Hollander, J.A., and Carelli, R.M. (2005). Abstinence from cocaine self-administration heightens neural encoding of goal-directed behaviors in the accumbens. Neuropsychopharmacology 30: 1464–1474.

Hursh, S.R. (1980). Economic concepts for the analysis of behavior. J. Exp. Anal. Behav. Econ. Concepts Anal. Behav. 34: 219–238.

Hursh, S.R., and Silberberg, A. (2008). Economic Demand and Essential Value. Psychol. Rev. 115: 186–198.

Izzo, A.A., Borrelli, F., Capasso, R., Marzo, V. Di, and Mechoulam, R. (2009). Non-psychotropic plant cannabinoids: new therapeutic opportunities from an ancient herb. Trends Pharmacol. Sci. 30: 515–27.

Kampman, K.M. (2019). The treatment of cocaine use disorder. Sci. Adv. 5: 1532–1548.

Kaplan, B.A., and Reed, D.D. (2014). Essential Value, Pmax, and Omax Calculator [spreadsheet application].

Lee, B.R., Ma, Y.-Y., Huang, Y.H., Wang, X., Otaka, M., Ishikawa, M., et al. (2013). Maturation of silent synapses in amygdala-accumbens projection contributes to incubation of cocaine craving. Nat. Neurosci. 16:.

Li, P., Wu, P., Xin, X., Fan, Y.-L., Wang, G.-B., Wang, F., et al. (2015). Incubation of alcohol craving during abstinence in patients with alcohol dependence. Addict. Biol. 20: 513–522.

Lu, L., Dempsey, J., Shaham, Y., and Hope, B.T. (2005a). Differential long-term neuroadaptations of glutamate receptors in the basolateral and central amygdala after withdrawal from cocaine self-administration in rats. J. Neurochem. 94: 161–168.

Lu, L., Grimm, J.W., Hope, B.T., and Shaham, Y. (2004). Incubation of cocaine craving after withdrawal: a review of preclinical data. Neuropharmacology 47: 214–226.

Lu, L., Hope, B.T., Dempsey, J., Liu, S.Y., Bessert, J.M., and Shaham, Y. (2005b). Central amygdala ERK signaling pathway is critical to incubation of cocaine craving. Nat. Neurosci. 8: 212–219.

Lu, L., Uejima, J.L., Gray, S.M., Bossert, J.M., and Shaham, Y. (2007). Systemic and Central Amygdala Injections of the mGluR2/3 Agonist LY379268 Attenuate the Expression of Incubation of Cocaine Craving. Biol. Psychiatry 61: 591–598.

Lujan, M.A., Alegre-Zurano, L., Martin-Sanchez, A., and Valverde, O. (2020). The effects of cannabidiol on cue- and stress-induced reinstatement of cocaine seeking behavior in mice are reverted by the CB1 receptor antagonist AM4113. BioRxiv Neurosci. 2020.01.23.916601.

Luján, M.Á., Cantacorps, L., and Valverde, O. (2019). The pharmacological reduction of hippocampal neurogenesis attenuates the protective effects of cannabidiol on cocaine voluntary intake. Addict. Biol. e12778.

Luján, M.Á., Castro-Zavala, A., Alegre-Zurano, L., and Valverde, O. (2018). Repeated Cannabidiol treatment reduces cocaine intake and modulates neural proliferation and CB1R expression in the mouse hippocampus. Neuropharmacology 143: 163–175.

Ma, Y.Y., Lee, B.R., Wang, X., Guo, C., Liu, L., Cui, R., et al. (2014). Bidirectional modulation of incubation of cocaine craving by silent synapse-based remodeling of prefrontal cortex to accumbens projections. Neuron 83: 1453–1467.

Mahmud, A., Gallant, S., Sedki, F., DCunha, T., and Shalev, U. (2016). Effects of an acute cannabidiol treatment on cocaine self-administration and cue-induced cocaine seeking in male rats. J. Psychopharmacol.

Mazzucchelli, C., Vantaggiato, C., Ciamei, A., Fasano, S., Pakhotin, P., Krezel, W., et al. (2002). Knockout of ERK1 MAP Kinase Enhances Synaptic Plasticity in the Striatum and Facilitates Striatal-Mediated Learning and Memory. Neuron 34: 807–820.

Nugent, A.L., Anderson, E.M., Larson, E.B., and Self, D.W. (2017). Incubation of cue-induced reinstatement of cocaine, but not sucrose, seeking in C57BL/6J mice. Pharmacol. Biochem. Behav. 159: 12–17.

Parvaz, M.A., Moeller, S.J., and Goldstein, R.Z. (2016). Incubation of Cue-Induced Craving in Adults Addicted to Cocaine Measured by Electroencephalography HHS Public Access. JAMA Psychiatry.

Pickens, C.L., Airavaara, M., Theberge, F., Fanous, S., Hope, B.T., and Shaham, Y. (2011). Neurobiology of the incubation of drug craving. Trends Neurosci. 34: 411–20.

Purgianto, A., Scheyer, A.F., Loweth, J.A., Ford, K.A., Tseng, K.Y., and Wolf, M.E. (2013). Different adaptations in AMPA receptor transmission in the nucleus accumbens after short vs long access cocaine self-administration regimens. Neuropsychopharmacology 38: 1789–1797.

Rodrigues, L.A., Caroba, M.E.S., Taba, F.K., Filev, R., and Gallassi, A.D. (2020). Evaluation of the potential use of cannabidiol in the treatment of cocaine use disorder: A systematic review. Pharmacol. Biochem. Behav. 196: 172982.

Roura-Martínez, D., Ucha, M., Orihuel, J., Ballesteros-Yáñez, I., Castillo, C.A., Marcos, A., et al. (2019). Central nucleus of the amygdala as a common substrate of the incubation of drug and natural reinforcer seeking. Addict. Biol.

Soria, G., Mendizábal, V., Touriño, C., Robledo, P., Ledent, C., Parmentier, M., et al. (2005). Lack of CB1 Cannabinoid Receptor Impairs Cocaine Self-Administration. Neuropsychopharmacology 30: 1670–1680.

Sun, A., Zhuang, D., Zhu, H., Lai, M., Chen, W., Liu, H., et al. (2015). Decrease of phosphorylated CREB and ERK in nucleus accumbens is associated with the incubation of heroin seeking induced by cues after withdrawal. Neurosci. Lett. 591: 166–170.

Terrier, J., Lüscher, C., and Pascoli, V. (2015). Cell-Type Specific Insertion of GluA2-Lacking AMPARs with Cocaine Exposure Leading to Sensitization, Cue-Induced Seeking, and Incubation of Craving. Neuropsychopharmacology 41: 1779–1789.

Trigo, J.M., Soliman, A., Quilty, L.C., Fischer, B., Rehm, J., Selby, P., et al. (2018). Nabiximols combined with motivational enhancement/cognitive behavioral therapy for the treatment of cannabis dependence: A pilot randomized clinical trial. PLoS One 13: e0190768.

United Nations publication (2020). World Drug Report. Wang, G., Shi, J., Chen, N., Xu, L., and Li, J. (2013). Effects of Length of Abstinence on Decision-Making and Craving in Methamphetamine Abusers. PLoS One 8: 68791.

West, E.A., Saddoris, M.P., Kerfoot, E.C., Carelli, R.M., and Hall, D. (2014). Prelimbic and infralimbic cortical regions differentially encode cocaine-associated stimuli and cocaine-seeking before and following abstinence. Eur J Neurosci 39: 1891–1902.

Whitfield, T.W., Shi, X., Sun, W.-L., and Mcginty, J.F. (2011). The Suppressive Effect of an Intra-Prefrontal Cortical Infusion of BDNF on Cocaine-Seeking Is Trk Receptor and Extracellular Signal-Regulated Protein Kinase Mitogen-Activated Protein Kinase Dependent.

Wolf, M.E. (2016). Synaptic mechanisms underlying persistent cocaine craving. Nat. Rev. Neurosci. 17: 351–365.

